# Cortical circuits modulate mouse social vocalizations

**DOI:** 10.1101/2022.05.20.492817

**Authors:** Benjamin Gan-Or, Michael London

## Abstract

Vocalizations provide a means of communication with high fidelity and information rate for many species. Male mice emit ultrasonic vocalizations (USVs) during female courtship. Diencephalon and brainstem neural circuits have been shown to control the production of USVs, however, the role of cortical circuits in this process is debatable. We show that male mice produced USVs following intracortical microstimulation (ICMS) in a specific location of their anterior cingulate cortex (ACC). Moreover, ACC Ca2+-imaging showed an increase in Ca2+ dynamics preceding USV initiation. Optogenetically suppressing ACC activity caused mice to emit fewer USVs during courtship. Neuropixel electrophysiological recordings in head-restrained male mice indicated a differential increase in neural activity in response to female social exposure (SE). The firing rate in SE trials where USVs were emitted was significantly higher when compared to SE leading to silence. Taken together, these results indicate that the ACC is a key node in the neuronal circuits controlling USV production.

## Introduction

Humans and a wide variety of animals use the vocal production of sound to facilitate a long-distance, high-throughput and high-fidelity means of communication that does not require visual or physical contact. Mice emit ultrasonic vocalizations (USVs) in various contexts (Chabout et al. 2015; Egnor and Seagraves 2016; Mun, Lipina, and Roder 2015; Peleh, Eltokhi, and Pitzer 2019). One prominent context in which mice emit simple and complex USVs is during courtship behavior (Holy and Guo 2005; McGill 1962; Ronald et al. 2020). Mouse USV emission is considered an innate behavior with little or no role for learning (Arriaga and Jarvis 2013; Hammerschmidt et al. 2012). Neural mechanisms in the subcortical forebrain and midbrain that directly control USV behavior were recently illuminated. It has been shown that nuclei in the preoptic area (POA), periaqueductal grey (PAG), and nucleus ambiguus are necessary and sufficient for USV emission and their modulation controls specific USV properties such as bout length or syllable amplitude (J. Chen et al. 2021; Hernandez-Miranda et al. 2017; Uwe Jürgens and Pratt 1979; Michael et al. 2020; Subramanian, Balnave, and Holstege 2021; Tschida et al. 2019). Naturally emitted USV bouts show a significant level of complexity (Hertz et al. 2020; Castellucci, Calbick, and McCormick 2018; Warren et al. 2021). At present, it is unclear whether POA and PAG induced USV bouts show the same degree of complexity. Other neural circuits, specifically cortical circuits and their projections, may contribute to the complexity of this behavior.

While in mammals such as humans, monkeys, and bats, there are cortical circuits dedicated to vocalization (U. Jürgens 2009), the involvement of the cortex in the production and control of USVs in mice is subject to debate. Arriaga et al., (Arriaga, Zhou, and Jarvis 2012) reported a direct projection from the primary motor cortex to laryngeal motoneurons. They also found that the cortex is required for USV repertoire modulation. In wild mice, the cortex was required to maintain USV pacing (Okobi et al. 2019). In sharp contrast, using genetic ablation of the entire population of excitatory neurons in the cerebral cortex, Hammerschmidt et al. (2015) have shown that the cortex is not necessary for USV production in mice, and that these mice lacking cortex from birth produce USVs indistinguishable from control mice. However, later analysis of these identical data (Ivanenko et al. 2020) showed that artificial deep neural networks could be trained to distinguish control mice USVs from those emitted by mice lacking cortex. Studies in mammals demonstrating the involvement of cortical circuits in vocalizations point to the anterior cingulate cortex (ACC) as a key cortical area. For example, intracortical electrical microstimulation (ICMS) of the ACC evokes vocalization in several mammals including rats, bats, guinea pigs, squirrel monkeys, macaques and humans (Bennett, Maier, and Brecht 2019; Coudé et al. 2011; Gooler and O’Neill 1987; Green et al. 2018; Müller-Preuss and Jürgens 1976; Roy, Zhao, and Wang 2016). Mice possess a homologue of some regions of the human ACC (Vogt et al. 2013); however, no study examined the ACC in relation to mice USVs. Furthermore, to our knowledge, no study has examined functional cortical neural activity in the mouse during USV production.

The evidence reviewed so far sets specific expectations for a possible role for cortical circuits in controlling USV emission. The sufficiency of subcortical circuits in USV production combined with the strong inhibition involved in this process indicates that a candidate cortical circuit is likely to play a modulatory role in the behavior. Hence, it is not expected that manipulating cortical circuits will be robust in initiating or abolishing USVs compared to manipulating subcortical circuits. Rather, it is expected that there will be aspects of cortical activity that would change prior to and in accordance with USV production.

To address this question of a cortical role in mice USV production, we have performed ICMS, fiber photometry recordings and optogenetic silencing of ACC. To further elucidate the neural activity correlate, we used electrophysiology in this region in vocalizing head-restrained mice to determine the specific cortical activity preceding USV production. We conclude that a sub-region of the mouse ACC is involved in the modulation of USV behavior during courtship.

## Results

### Intracortical Microstimulation in Anterior Cingulate Cortex Triggers Ultrasonic Vocalizations

In view of the seemingly contradictory evidence for the involvement of the cortex in the control of USVs in mice (Arriaga and Jarvis 2013; Hammerschmidt et al. 2015), we have tested if electrical activation of the mouse’s cortex can trigger USVs. Previous studies have demonstrated that ICMS in the cortex of various other mammals triggers vocalizations (Bennett, Maier, and Brecht 2019; Green et al. 2018; U. Jürgens 1995; Okobi et al. 2019). Using ICMS, we performed a systematic mapping of the motor and premotor areas and tested if USVs can be induced in male mice. Male mice only emit USVs while awake, therefore a systematic large-scale mapping of possible vocal nodes requires head-restraint. Male mice naive to female exposure were habituated to head restraint (Fig. 1A). To prevent any hint of female scent in the ICMS experiments, mice were never exposed to females on the head restraint apparatus and females were never brought into the rig throughout the habituation procedure (Fig. 1B). No mice produced any detectable spontaneous USVs while head-restrained (n=23 mice, total recording time = 61 hours (see also Weiner et al., 2016). This complete absence of spontaneous USVs provided an ideal setting to test for USVs triggered by ICMS.

**Figure 1:**
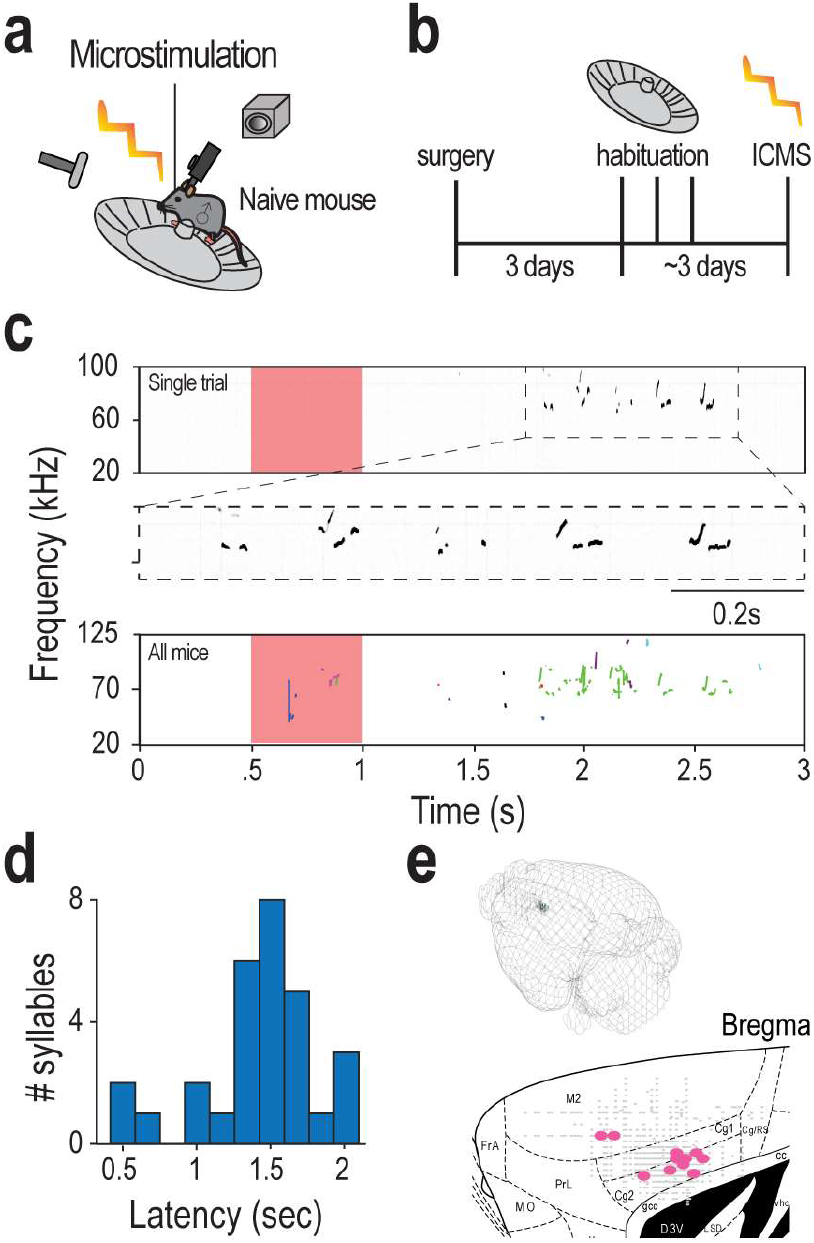
Microstimulation in a specific location in anterior cingulate cortex induces ultrasonic vocalizations. **A**. Schematic illustration of the experimental setup. A glass pipette is used to induce microstimulation in the cortex of a head-restrained male mouse positioned on a running wheel, while an ultrasonic microphone records USVs and a camera records any behavioral response. Male mice are naïve to female exposure from weaning at P21. **B**. Schematic illustration of experimental preparation. After head-restraint, mice recover for three days and subsequently undergo habituation to the head-restraint apparatus lasting about three days. **C**. A single stimulation trial spectrogram example of USVs elicited by the mouse following ACC microstimulation (stimulus duration of 500ms). Below is a zoomed-in spectrogram showing that the evoked sound is complex and in the ultrasonic range. Further below is an overlay of all USVs (n=29 syllables) induced in all mice (n=8 mice). **D**. Histogram of latency from stimulation onset to USV onset. **E**. A 3D visualization of all stimulation sites, with USV-inducing sites in green and quiet sites in grey. Below is a sagittal section with coordinates of locations tested by ICMS (gray markers denote locations that did not induce any USVs, while pink markers denote locations where at least one USV was induced).

We then performed penetrations over the cortical area at various depths at each location (advancing in vertical steps of 50 µm). Following the work of Graziano (Graziano, Taylor, and Moore 2002, 200), in which long duration ICMS resulted in complex naturalistic movements in monkeys, we have used at each location both short and long duration stimulation trains (short: 50 ms, long: 200-500 ms, Methods). In five mice, short- and long-duration ICMS in the primary motor cortex (M1) and secondary motor cortex (M2) produced a large variety of simple (twitches) and complex movements (see also (Hira et al. 2015)) but have not resulted in any USV production (# USVs =0, 1,148 penetrations, 11,203 stimulations). In addition, we have stimulated at the coordinates reported in a previous report (Arriaga and Jarvis 2013) within M1 but failed to induce USVs. Then, we set out to test ICMS in the cingulate cortex (ACC) (Vogt and Paxinos 2014) and in sharp contrast, at a clustered location in the anterior ventral part of the ACC, long-duration ICMS evoked USVs (Bregma: 700±100 µm medial 800±300 µm anterior 1500±300 µm depth Fig. 1E). An example of an individual USV and several USVs in succession following a 500ms long ICMS is depicted in Fig. 1C. Importantly, these vocalizations were in the ultrasonic range and did not represent stress-related sonic vocalizations (Grimsley et al. 2016). An overlay of all USVs followed all ICMS events (in successful stimulation sites) across all mice is shown in Fig. 1C (total ICMS sites= 1974, successful sites= 12, number of triggered USVs = 29, n=8 mice). Only long-duration stimulations resulted in USVs, while short-duration ICMS at the same locations were never followed by USVs. Long-duration ICMS-induced USVs had an average frequency of 81kHz and varied in duration (see full statistics in Table S1). These characteristics are not significantly different from typical USVs emitted during courtship by head restrained male mice (Weiner et al. 2016). USVs were only triggered by long-duration ICMS with relatively long latencies to USV emission (average latency to first USV: 1.5 ± 0.7 s, N=8, Fig. 1D), suggesting a complex involvement of ACC in ultrasonic vocalization. This response latency is in agreement with ICMS-induced USVs in the ACC of monkeys and rats (Bennett, Maier, and Brecht 2019; Uwe Jürgens and Ploog 1970). In conclusion, we have identified a location within the ACC of male mice where long-duration ICMS evokes the initiation of USVs, similar to courtship USVs, albeit with a low success rate and long latency.

### Neural Activity in the Anterior Cingulate Cortex Precedes Vocal Onset

To explore the activity of the ACC neuronal population during natural courtship USV behavior, we used fiber-photometry population Ca^2+^-imaging (Stroh et al. 2013). We performed stereotactic injections of AAV viral vectors encoding for Ca^2+^ indicator AAV9.cag.GCaMP6s (see Methods)(T.-W. Chen et al. 2013) into the location identified using ICMS in the left ACC. (Fig. 2A). After expression, an optical fiber (diameter = 200 μm, NA = 0.22) was implanted in the coordinates specified above and we measured Ca^2+^ dynamics using fiber photometry in a natural courtship setting as depicted in Fig. 2B (see Methods). Mice implanted with an optical fiber were placed in a fresh cage for a short habituation period (5 minutes). Then the session began and for a duration of 20 minutes a female mouse was repeatedly presented to induce USV production and then immediately removed from the cage. During the session, the Ca^2+^ signal and microphone signal were recorded continuously. The male mice often responded to female entry by emitting USVs. We defined a song as a succession of syllables with a surrounding silent period of at least 10 seconds of quiet (see methods for syllable and song definitions) (Holy and Guo 2005). Six mice were recorded for at least one session, totaling 33 hours of recording with 2,176 songs (mean: 363 songs per mouse).

**Figure 2:**
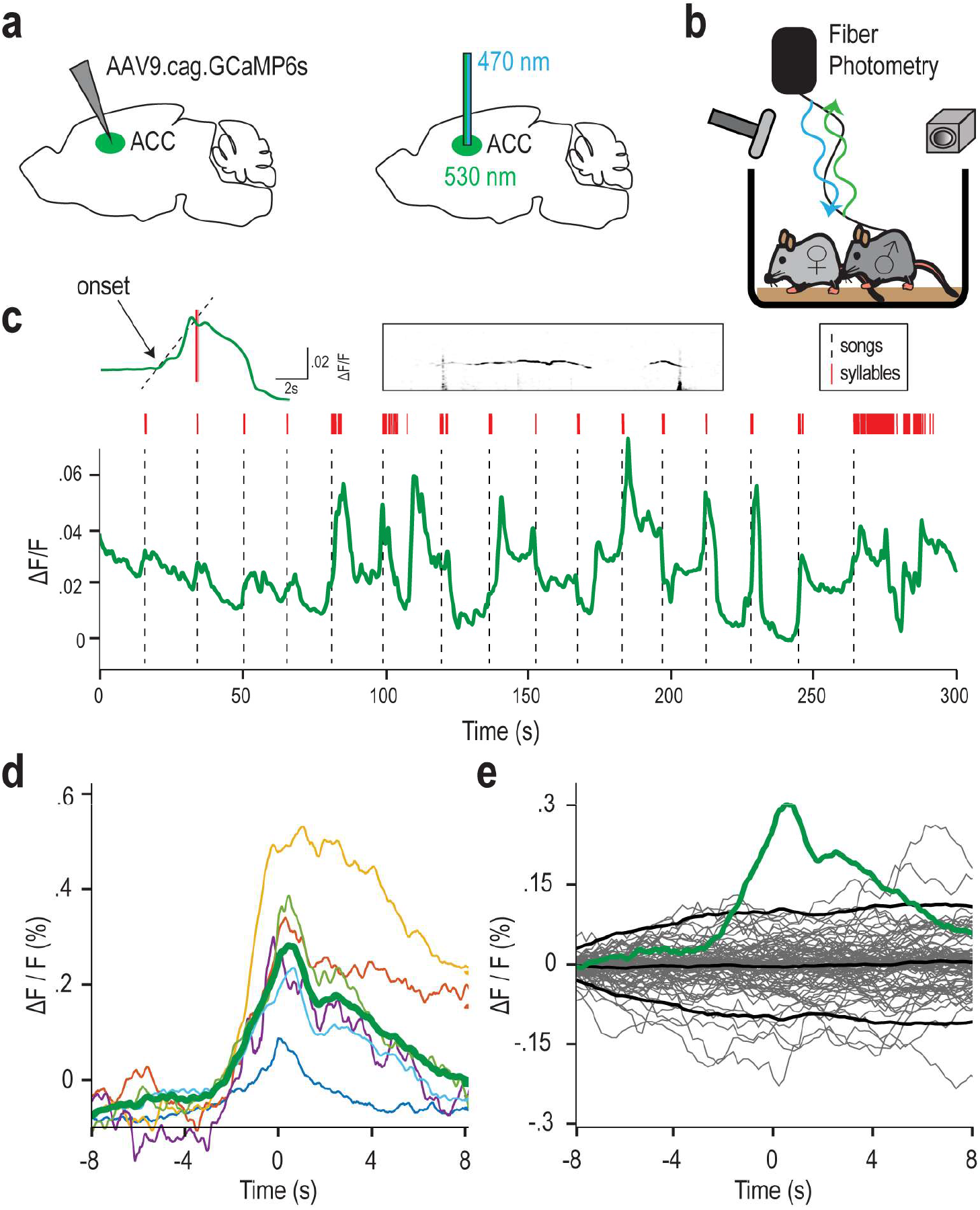
Anterior Cingulate Cortex Ca^2+^ activity correlates with song initiation. **A**. Virus with Ca^2+^ indicator was injected two to three weeks before recording. Recordings were made from the left ACC using blue light to activate the Ca^2+^ indicator and green light was detected in the photodiode. **B**. Schematic illustration of fiber photometry setup. Male mice with fiber implants were allowed to move freely in a cage, while a female was introduced to elicit USVs. **C**. Example session of 5 minutes in duration. The green line represents Ca^2+^ signal, thin red lines are syllables, and the dotted black line represents song initiations. Above is zoom-in on a single song (with a dotted line showing activity onset calculation), with the spectrogram to the right showing the first two syllables in the song. **D**. Average Ca^2+^ signals for all mice, triggered on the first syllable in a song separated by at least 10 seconds. **E**: Permutation test trials using random trial times plotted in gray. The average from D in green is plotted against the average and ±2 SD of the permutation test average in black.

Ca^2+^ transients showed a clear temporal correlation with song initiation (Fig. 2C). Averaging the Ca^2+^ signal aligned on song initiation (the time of occurrence of the first syllable in every song) revealed an elevation in Ca^2+^ activity (Fig. 2D). To test whether this elevation in the Ca^2+^ signal is significant, we used a permutation test. We generated a sequence of random times to trigger the Ca^2+^ signal and calculated the distribution of the expected Ca^2+^ signal from the data (Fig. 2E). We then compared the actual signal to the expected one for each mouse. 5 out of 6 mice passed this test with p<0.05. To test whether the Ca^2+^ signal preceded or followed the song onset, we defined the Ca^2+^ onset by fitting a straight line to the range of 10%-90% of its peak amplitude and extrapolated to zero-crossing (see Fig. 2C upper-left inset). We found that the Ca^2+^ elevation begins prior to song onset. The average latency from Ca^2+^ signal onset to song initiation was 2.3±3.8 sec (n=6). This temporal difference is in good agreement with the delay reported in the ICMS experiment above (taking into account the slow dynamics of the Ca^2+^ signal and its low pass filtering). These results demonstrate an elevation in activity in the ACC location discovered via ICMS. This elevation occurs prior to song initiation while mice are engaged in natural courtship behavior, further indicating cortical involvement in this behavior.

### Silencing Anterior Cingulate Cortex Neural Activity Reduces Syllable Production and Song Initiation

Previous studies have shown that activation of subcortical regions such as the PAG and POA is sufficient for triggering USVs and controlling some of their basic properties (such as syllable volume and bout duration) (J. Chen et al. 2021; Gao et al. 2019; Michael et al. 2020; Tschida et al. 2019). Nevertheless, the effect of manipulation of cortical activity during courtship on vocalization has not been tested. In order to test if silencing of the ACC would affect courtship USVs, we expressed the optogenetic silencing actuator *Guillardia* theta anion-conducting channelrhodopsin (GtACR2) (Malyshev et al. 2017, 2) unilaterally in the left hemisphere ACC of male mice to suppress neural activity using light (Fig. 3A). After recovery from optical fiber implantation, mice were placed in a clear cage while connected to the laser without light stimulation while USVs were recorded continuously. The session started with 5-minute habituation without a female. Afterward, a female was introduced into the arena for an additional 20 minutes (experimental schematic in Fig. 3B). After two days, mice again met a female for 20 minutes with the laser-activated for 50 interspersed stimulation trains (duration = 10 seconds, frequency = 20Hz, pulse duration=10ms). Example segments from the light sessions demonstrating eight stimulation trains are depicted in Fig. 3C. The final post-stimulus session was conducted with the same behavioral protocol after two days. To test if light stimulation affected USVs, we compared the number of USVs emitted during light-delivery periods (blue sections in Fig. 3C) versus concurrent time intervals in the absence of light stimulation. The stimulation did not produce any observable disruption of normal courtship behavior, nor did it prevent vocal behavior. However, optogenetic suppression of ACC activity reduced the number of syllables produced, whereas the syllable main frequency component remained normal (Fig. 3C). Relative to baseline, the number of syllables emitted in the light delivery sessions was reduced and returned to baseline in the post-stimulus session (Fig. 3D; baseline: 1007 ± 174, suppressed: 343 ± 67, post: 681 ± 92, p < 0.05, N=5). Mice also initiated fewer songs during ACC suppression compared to the baseline session, and here too, recovery occurred in the post-stimulus session (Fig. 3E; baseline: 55 ± 3, suppressed: 48 ± 3, post: 57 ± 2, p<0.05, N=5). A control group of mice injected with AAVdj.CMV.eGFP were tested using the same protocol. In the control group, no significant changes were found in the number of emitted syllables during stimulation trains or in song initiations (Fig. 3E,D, # syllables: baseline: 801 ± 159, suppressed: 767 ± 135, post: 964 ± 198, # songs: baseline: 54 ± 6, suppressed: 58 ± 5, post: 60 ± 4, p >0.05, N=6).

**Figure 3:**
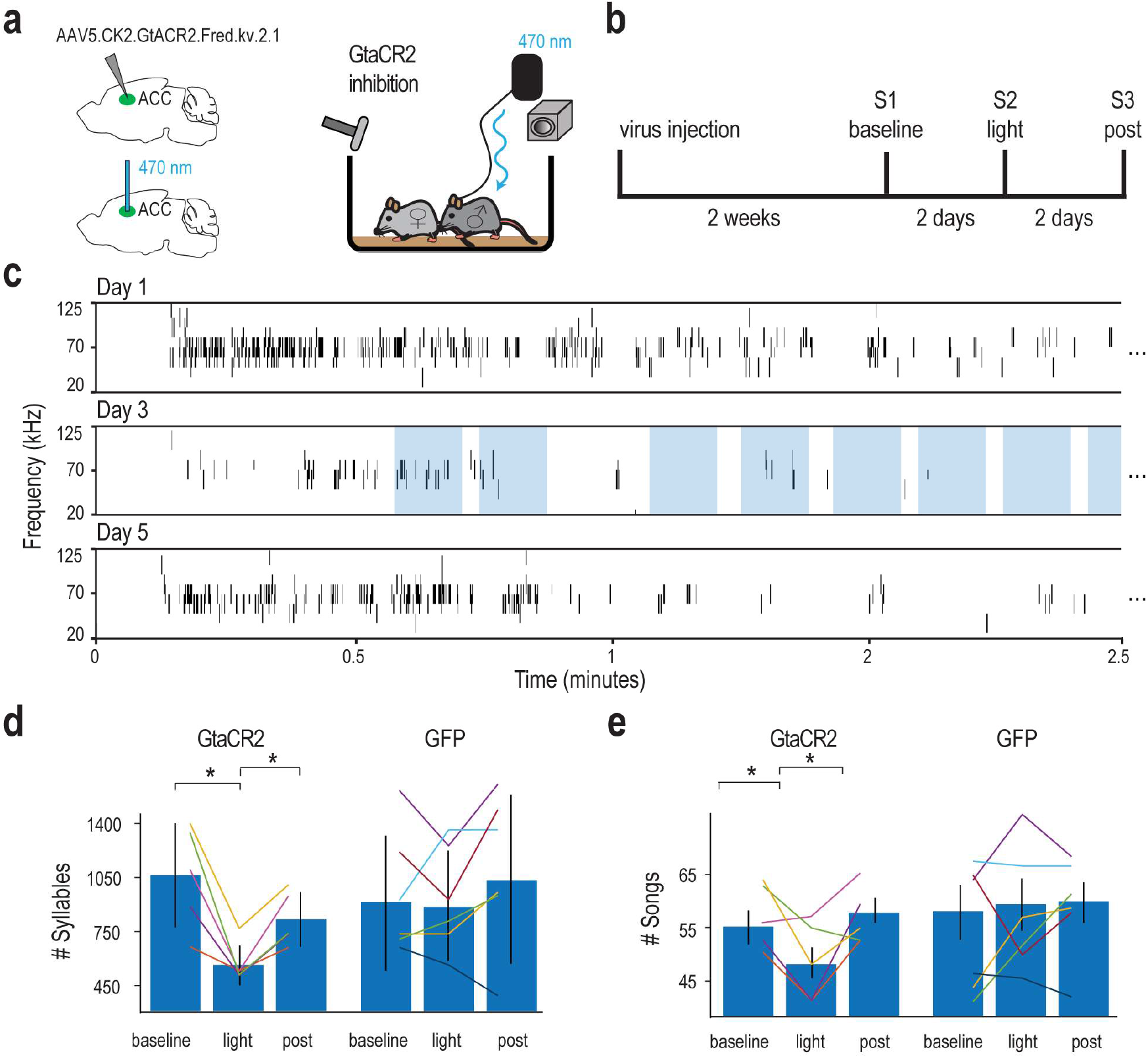
Suppressing anterior cingulate cortex activity reduces emitted syllables and song initiation. **A**. Schematic illustration of optical stimulation setup. A male mouse with optical fiber implant and injected GtACR2 inhibitory virus freely moves in a cage, while a female is placed by hand into the cage during experimental trials. **B**. Schematic of the experimental timeline. Two weeks after virus injection, to allow for strong expression, session 1 baseline USV performance is assessed. After two days, session 2 USV emission is assessed with simultaneous optogenetic silencing. In session 3, after two days, post-stimulus USVs are assessed for recovery. **C**. Example 2.5 minutes of USVs from a baseline session trial, stimulation session (light) trial, and post-stimulus session. Individual stimulation epochs are shown in light blue during the light trial. Mice made fewer syllables in the stimulation (light) sessions. **D**. Mice emitted fewer syllables during ACC suppression. Percentage of syllables across the different sessions with average in blue bars with standard error. Control (GFP) sessions are compared on the right side. Baseline mice syllable emission is variable, but the stimulation session showed a significant decrease in comparison to the post-stimulus session two days after the stimulus session. **E**. Mice emitted fewer songs during ACC suppression. The number of songs across the different sessions with average in blue bars with standard error. Control (GPF) sessions are compared on the right side. Compared to baseline, mice emitted fewer songs during the stimulation session and fully recovered in the post-stimulus session.

The results of these experiments suggest that a reduction in ACC neural activity is associated with a reduction in syllable emission and song initiation, which further strengthens the conclusion that the ACC is an important node in the USV-controlling network.

### Differential Responses of Anterior Cingulate Cortex and Secondary Motor Cortex Neurons to Social Exposure Conditioned on Song Initiation

The series of experiments described above provides one correlative and two independent causal lines of evidence showing that the ACC is involved in USV control. Despite this, the recorded signals and manipulations do not permit spatial and temporal resolution down to the level of individual neurons and spikes. We next wanted to explore the role of individual neurons in this region of the ACC during courtship USVs. We, therefore, performed electrophysiological recordings at a greater spatial and temporal resolution. Adult male mice were habituated to the head-restraint apparatus used in Fig. 1A. We then used a Neuropixel system (Jun et al. 2017), see Methods) to record at a depth of 2.5-3 mm in the cingulate cortex (ACC) and secondary motor cortex (M2). USVs and video were simultaneously recorded (Fig. 4A). To trigger USVs during the recording, female mice were repeatedly introduced by an experimenter. Mice usually responded to female presentation by emitting songs (65% of female presentations resulted in songs in 1079 of 1661 presentations). In contrast to the ICMS experiment, here male mice were head-restrained while exposed to females. As a result, in some cases, this led to songs indirectly induced by female presentation (constituting 11% of total song production).

**Figure 4:**
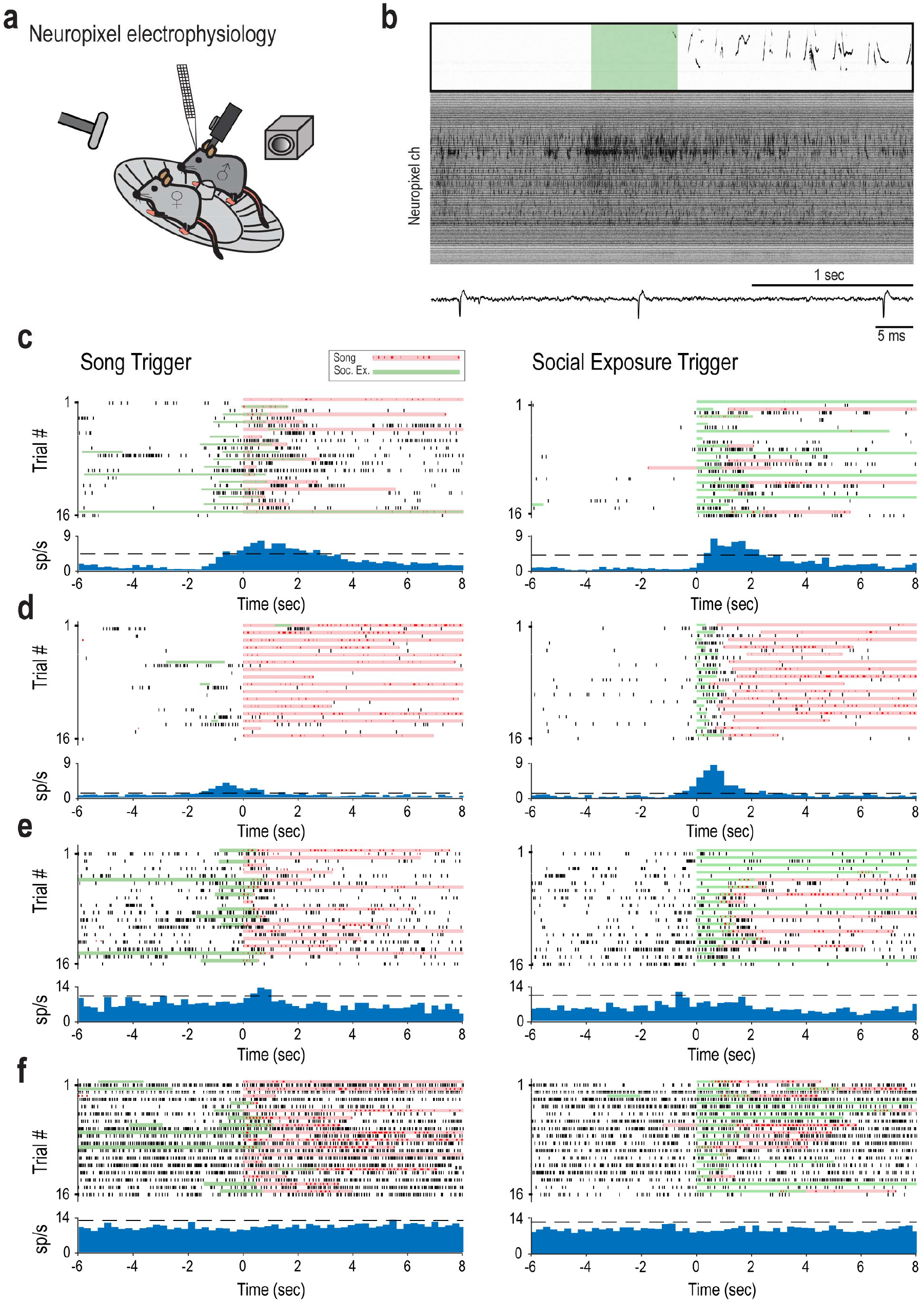
Both social exposure and song initiation modulate single unit activity. **A**. Schematic illustration of the experimental setup. A Neuropixel probe is inserted into the ACC of a head-restrained male mouse (positioned on a running wheel), while an ultrasonic microphone records USVs and a camera records any behavioral response. **B**. Example of the recording quality of all channels in response to social exposure and subsequent song initiation. Social exposure epoch is in light green superimposed over the spectrogram of the recorded sound. Below is a zoomed-in example of a single channel. Only units passing 10 standard deviations above the mean waveform value are included in the analysis. **C-F**. Example of single units triggered on both song and social exposure. **C**. This example shows a single unit responding to social exposure and song. **D**. Here, a single unit responds more strongly to social exposure as compared to song initiation. **E**. A unit with song initiation inducing a weak response, but social exposure does not. **F**. An example of a single unit responding to neither song nor social exposure.

Electrical recordings had a high signal-to-noise ratio, obviating the need for spike sorting and allowing us to focus on single unit activity (Fig. 4B, see methods). In total, we recorded 130 units in 3 mice. We use the term social exposure (SE) to denote the female presentation as the male was not always directly interacting with the female. Figure 4C-F shows example raster plots and Peri-event time histograms (PETHs) triggered on song initiation or SE. Each row shows a different single unit. We found a variety of units responding to social exposure (Fig. 4D), song initiation (Fig. 4E), both (Fig. 4C) or none (Fig. 4F).

PETHs for all 130 units show that the population of neurons responded both to SE and song initiation (Fig. S4A,B). The time course of the population PETH triggered on song initiation is in good agreement with the average Ca^2+^-signal response shown in Fig. 2E. To quantify the modulation of an individual unit’s activity with respect to social exposure and song initiation, we selected units if their activity deviated by two or more standard deviations compared to baseline (determined by a permutation test). We found 34 units that passed this test and showed a significant change in firing rate with respect to either SE or song initiation. Both M2 and ACC had equal amounts of responsive neurons (Fig. 5A).

**Figure 5:**
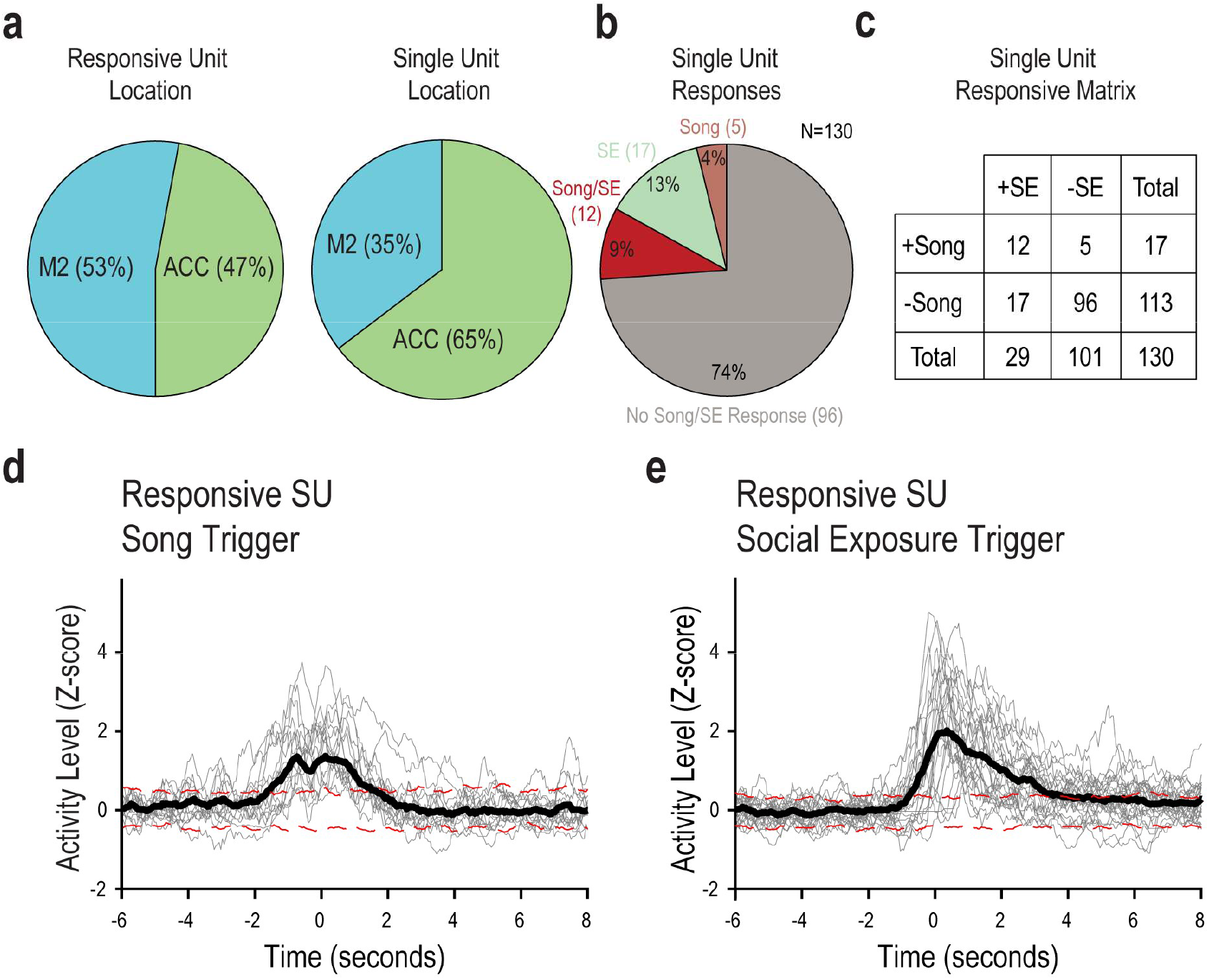
Two overlapping populations of neurons modulate their activity preceding song initiation and social exposure. **A**. Basic response properties across all single units. The majority of single units recorded were in ACC, yet out of the 34 (22%) responsive single units, only 50% were in ACC. **B**. 25% of neurons recorded were responding to either song or social exposure. **C**. The responsive matrix lists the number in each group. +Song represents the neurons responsive to song, while +SE represents neurons responsive to social exposure. These categories do not indicate the absence of one of the groups (i.e., -SE only indicates lack of response, not lack of social exposures in that group). **D**. Averaging all 34 responsive single units on social exposure reveals a strong response centered around 1 second after social exposure in the PETH as compared to ±2 SD of a permutation test. This response is stronger than when averaging on song initiation. By separating trials when song or quiet follows social exposure, we can explore a USV modulatory effect. **E**. For 17 song responsive neurons, averaging the PETH on song initiation yielded a significant increase in activity preceding song initiation as compared to the dotted red lines representing ±2 SD in a permutation test. However, most song initiations were preceded by social exposures, limiting the scope of this result.

The average PETHs of only social exposure responsive (SE+) single units show an elevation in firing rate when compared to a permutation test (Fig. 5B,C). The PETH with respect to song initiation (S+) reveals song-related modulatory activity in half (17/34) of responding units. Thus, the average song-PETH shows a significant firing rate elevation beginning at 1.6 seconds prior to song (Fig. 5D). About one third of the units (36%, 12/34) show a significant elevation in firing rate when triggered on either SE or song initiation (examples in Fig. 4). In our experimental paradigm, it was impossible to determine the exact moment of social exposure and, therefore, difficult to draw conclusions about the fine temporal relationship between SE and the change in firing rate. Nevertheless, we can conclude that 85% (29/34) of responding neurons have demonstrated a significant elevation in firing rate in response to SE (Fig. 5E).

Figure 5 suggests that the average firing rate of some units in the ACC and in M2 is increased prior to song initiation. However, it is difficult to determine if the elevation indeed reflects involvement in song emission because the song initiations were almost always preceded by social exposure to a female stimulus. In this experiment, 89% of song initiations were preceded by a social exposure (Fig. 6A). Only 11% of songs were initiated in isolation. Therefore, it is possible that the elevation in firing rate seen in the PETH of S+ units only reflects a response to SE. We examined the latency from social exposure to vocal production and found that most songs occur within 2 seconds of social exposure (Fig. 6B); therefore, it is impossible to separate the two responses fully.

**Figure 6:**
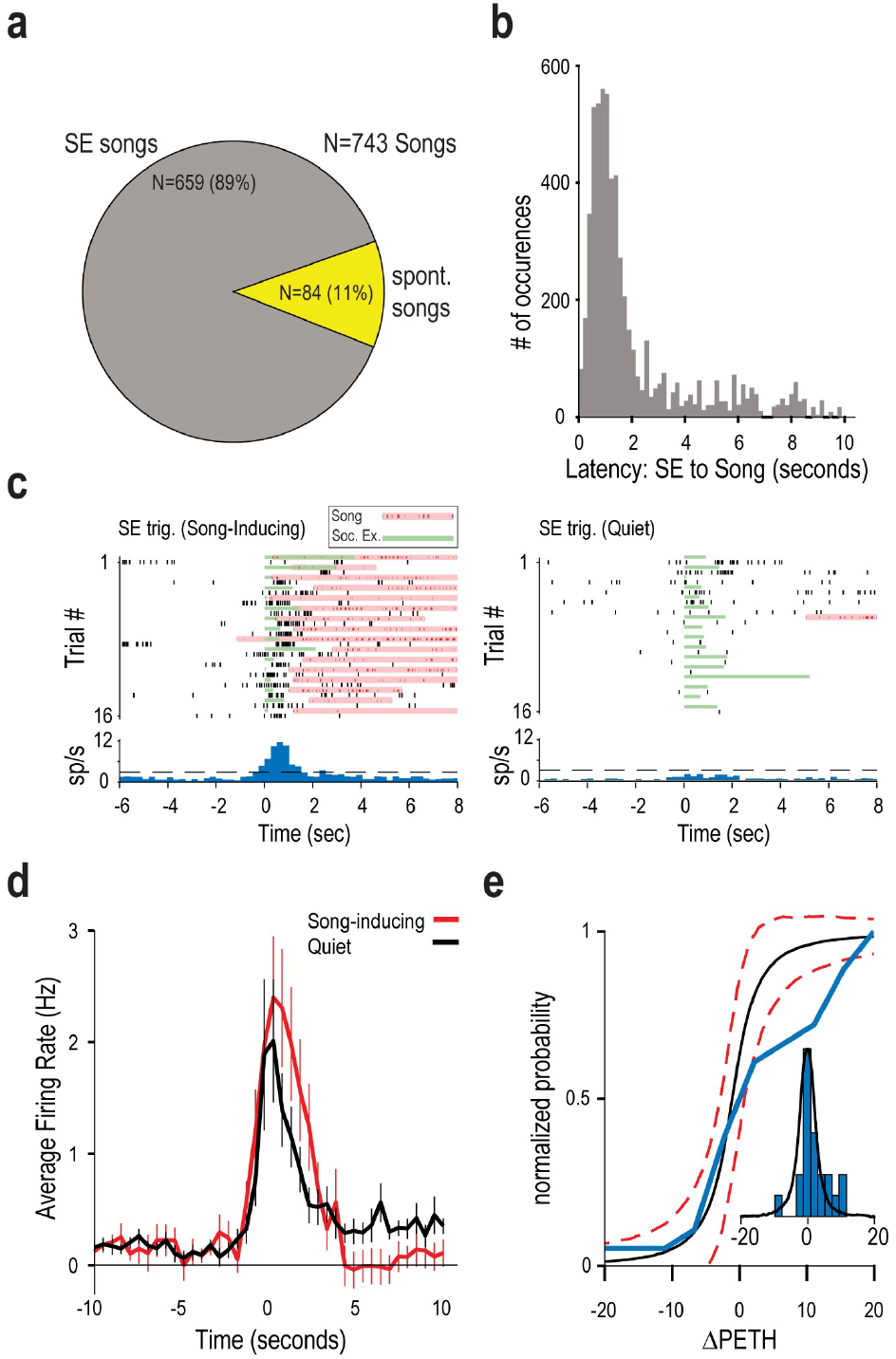
Conditioning on social exposure reveals an ultrasonic vocalization modulatory effect on anterior cingulate cortex and secondary motor area neural activity. **A**. Majority of songs are preceded by social exposure. In 89% of song emission, mice were exposed to a female, while only 11% of songs were initiated in isolation (at least 5 seconds of isolation from female SE). **B**. Most songs occur ∼1-2 seconds following social exposure. Thus, social exposure is a confound when averaging neural activity triggered on song initiation. **C**. Two example PETHs from a single unit showing responses triggered on SE, separated to SE resulting in a song or not (quiet). In the above PETH, a Song-Inducing SE reveals an increase in activity about 1 second after exposure. On the other hand, a Quiet social exposure shows no increase in activity. **D**. Average SE triggered PETH across all responding neurons (N=18, maximum 40 social exposures per neuron) split between Song-Inducing (red) and Quiet (black) social exposures. Song-Inducing social exposures have a higher firing rate following exposure and a subsequent inhibition. For each unit, the SE-triggered PETH for song-inducing and quiet conditions were calculated (Fig. 6S) and the difference between them was measured as ΔPETH. **E**. The cumulative distribution of ΔPETH is shown (blue). The black line (predicted) is calculated by a permutation test. 1000 randomly shuffled groupings of the Quiet/Song-Inducing signal were chosen and the ΔPETH was averaged. The ΔPETH distribution (blue) is shifted to the right compared to the random grouping, representing the increase in single units preferring Song-Inducing social exposures.

However, not every SE resulted in song emission (1079 out of 1661 SE resulted in songs (65%)). Therefore, we have computed the SE-triggered PETH separately for SE that were followed by a song and for the SE followed by silence. Fig. 6C depicts PETHs of one SE+ single unit, separating out the trials with songs or silence following social exposure. Clearly, this unit shows a song-inducing correlation. Out of 34 responding units, we selected 18 neurons with an average firing rate greater than 1Hz during social exposure for this analysis. We further categorize their responses as Song-Inducing (10/18), Quiet (3/18), or Agnostic (5/18) single units (the individual PETH of all units are depicted in figure S6). The figure shows that the majority of SE+ units modulate their SE triggered activity depending on whether the mouse will produce a song or remain quiet. Therefore, even considering the confound of song-triggered PETHs, which could reflect SE response, ACC neurons modulate their activity with respect to song initiation and their modulation has predictive power.

By averaging all SE-triggered PETHs separately for song trials and quiet trials, we found that neurons in sum modulate positively their firing rate in response to Song-Inducing social exposure (Fig. 6D and see also Fig. S6). In agreement with the average elevation in firing rate, analysis of the responses of individual neurons shows the same tendency. We quantified this by analyzing the distribution of individual cells using the difference between the PETH in the Song-Inducing and Quiet conditions (ΔPETH) across all responsive single units. We found that the real distribution of ΔPETH is shifted towards single units responding more towards Song-Inducing preference in comparison to a shuffled distribution (Fig. 6E). Thus, vocal production modulates the social exposure response of ACC and M2 neurons.

## Discussion

We demonstrated that the cortex plays a role in the male mice USV courtship behavior. Intracortical microstimulation in a restricted area of the ACC was sufficient to invoke USVs in head restrained mice that were absolutely mute otherwise. This was achieved in male mice who never encountered a female mouse or received any other stimulation such as female mouse urine or other odorants. Furthermore, successful trials, in which USVs were elicited, occurred only with long-duration stimulation in a restricted area of the ACC and not in other candidate cortical areas. Using fiber photometry, we identified Ca^2+^ signals from the same ACC location which precede song initiation. This elucidated a possible function for ACC in initiating songs. We also showed that when neural activity in the ACC is suppressed optogenetically (unilaterally), mice produce fewer syllables and initiate fewer songs in the presence of female mice. These results propose the ACC as an important node in the USV-controlling network. Lastly, using electrophysiology, we have shown that units in M2 and the ACC respond with firing rate elevation to female exposure and prior to song initiation. Moreover, many units respond to female exposure differently depending on the future vocal response, whether that response is a song or a quiet period.

Our results fit well with the literature on the involvement of the cortex in vocalizations in many mammals such as monkey, rat and bat (Bennett, Maier, and Brecht 2019; Gavrilov, Hage, and Nieder 2017; Rose et al. 2021). Purposeful, but not spontaneous, vocal behavior in monkeys is associated with increased ACC activity. In bats, prefrontal neurons fire differentially in response to calls from different individuals in a social setting. Previous ICMS and local field potential recording studies in rats demonstrated a role for the ACC in USV production (Bennett, Maier, and Brecht 2019; Burgdorf et al. 2007; Saito and Okanoya 2017). Rats produce a type of 50kHz USV, which is induced spontaneously or under a variety of conditions from handling to feeding (Laplagne and Elías Costa 2016). Tickling or ICMS of somatosensory areas also evokes this USV behavior. (Burgdorf, Panksepp, and Moskal 2011; Harmon et al. 2008; Ishiyama and Brecht 2016). Rat USVs may be a general expression of emotional state, whereas mice USVs are likely directed towards certain well-defined behaviors as in courtship. Interestingly, in a strain of wild mice (Okobi et al., 2019), vocalizations arise in a unique behavioral context, and stimulation in the motor cortex disrupts vocalization timing. In subcortical areas, recent results reveal the direct role of the POA, PAG and nucleus ambiguus in USV production (J. Chen et al. 2021; Hernandez-Miranda et al. 2017; Michael et al. 2020; Tschida et al. 2019). The short latency and greater reliability of evoked USVs in those studies, as compared to USVs triggered by ICMS in the ACC, demonstrates the direct connection between subcortical brain areas and USVs. However, the role of the ACC in affecting these areas has not been studied in the mouse. Moreover, the long latency and the difficulty in inducing USVs could also reflect the implications of tonic inhibition present in subcortical regions controlling USVs.

Importantly, activating the cortex with ICMS may induce non-specific neural activity in remote areas due to stimulation of passing-by axons, and therefore it is difficult to rely on ICMS for localization of function (Histed, Bonin, and Reid 2009). However, here we support this localization with three additional lines of evidence. The fiber photometry and electrophysiological recordings show specific activity in a significant population of neurons with respect to song emission. The optogenetic silencing of ACC, which leads to a reduction in syllables and songs, further suggests that ACC takes an important role in the USV-controlling network.

Our results challenge previous studies focusing on the anatomical aspects of courtship USVs. A previous finding showed that the motor cortex in mice has a direct monosynaptic connection to the nucleus ambiguus, where laryngeal motor neurons reside (Arriaga, Zhou, and Jarvis 2012). However, our electrical stimulation in the motor cortex, while successful in eliciting complex motor responses, failed to elicit any vocal response. These two results could still be consistent if, for example, other areas are continuously inhibiting vocal production. Indeed, previous work revealed that POA inhibition onto PAG inhibitory interneurons evokes USV production. (J. Chen et al. 2021). However, this putative PAG “gate” must either spontaneously fire or receive input in order to trigger USVs when inhibition is removed. Alternatively, vocal production could require coordination of several processes, and the motor cortex might only exert partial control. In contrast to our motor cortex ICMS, our anterior cingulate cortex ICMS successfully triggered USVs. This suggests that the ACC resides higher in the control hierarchy of vocal control. When the cortex (including the ACC) is removed through genetic manipulation, mice still vocalize (Hammerschmidt et al. 2015). Since the analysis showed no statistical differences between the USVs produced by control and mutant mice, the authors concluded that the cortex is not necessary for USV production and control. However, recent work from the same authors has instead shown that a more sophisticated analysis can separate USVs produced by controls and mutants (Ivanenko et al. 2020). Therefore, our results are in contrast to the earlier finding but are consistent with the latter and with the view that the cerebral cortex is likely to exert complex control over innate behaviors.

The involvement of the ACC in the mouse USV process opens the option that this process is more complex and plastic than currently assumed. It is already known that mice vocalize differently when social context is varied (Chabout et al. 2012; 2015; Ey, Chaumont, and Bourgeron 2020; Sasaki, Tomita, and Kanno 2020), and that they can modulate their pitch in a cross-strain fostering scenario (Arriaga and Jarvis 2013). The access in the mouse to specific cell populations and genetic manipulations provide a unique opportunity to explore and manipulate higher-order control on the USV-controlling network and ultimately relate it to human speech in health and in various forms of neurological and genetic conditions in which it is disrupted (Ciucci et al. 2010; French et al. 2019; Grant et al. 2014, 201; Michetti 2012, 20; 2012, 20; Offen et al. 2018; Scattoni et al. 2008; Selimbeyoglu et al. 2017; Shu et al. 2005; Wöhr 2014; Young et al. 2010, 201).

## Materials and Methods

### Animals

We used C57BL/6 male mice (at least 8 weeks old). The Hebrew University Animal Care and Use Committee approved all experiments. IACUC NS-16-14216-3 “Functional identification of brain regions involved in ultrasonic vocalization in mice.”

### Surgery and viral vectors injections

During surgery, mice were anesthetized with isoflurane (1-2% by volume in O2 LEI medical). Rymadil analgesia (10mg/kg body weight, 200µl injection volume) was administered prior to incision. Mice were put in a stereotactic frame (Narishige, Japan/Luigs & Neumann GmBH) and a small craniotomy <1 mm in diameter was made over the left anterior cingulate cortex (dura was not removed). Virus-containing solution was slowly injected (1-2 nl/sec) through a quartz pipette (pulled on P2000, Sutter instruments,) using a Nanoject 3 system (Drummond Scientific Company, PA) (500 nl, one site per animal at a depth of 1400-1700 um). After injection, the craniotomy was sealed with bone wax and the skin was attached with VetBond (3M). Two to three weeks after the virus injection, the mice were prepared for optical stimulation. Surgery was performed under isoflurane anesthesia and Rymadil analgesia was given. For the optical stimulation experiments, a custom-made fiber optical cannula was subsequently implanted over the injection site. For animals undergoing fiber photometry experiments, an optical fiber was directly connected to the skull with dental cement under the same conditions. For ICMS experiments, a head post was fixed on the skull under the same surgical conditions, and subsequently awakened and allowed to rest for 2-3 days before ICMS began. For electrophysiology recordings, a head post and ground pin were fixed on the skull under the surgical conditions, and subsequently awakened and allowed to rest for 2-3 days before recording sessions began.

### Sound recording and USV analysis

In all experiments, sound was recorded to capture USVs using an ultrasonic microphone (Avisoft, CM16/CMPA) amplified (either by CMPA40-5V or UltraSoundGate 116H) and digitized at 250kHz (either by National Instruments Card or Avisoft RECORDER). In fiber-photometry and optogenetic experiments, males were isolated for 5 minutes in a new cage and presented with a female either by adding it to the same cage or by presenting it on the experimenter’s hand. In electrophysiology experiments, head-restrained males were presented with a female mouse or female mouse urine only on the experimenter’s hand to induce vocal production. We parsed USVs using an algorithm previously published available at https://github.com/london-lab/MouseUSVs/tree/master/USV_Parsing (Hertz et al. 2020). In brief, the raw audio signal is cleaned for common noise sources and pixels in the spectrogram are removed if they are isolated from other pixels. For the algorithm parameters, we used a frequency cut-off between 20-125kHz and a square size of 3. In all subsequent analysis, a syllable is defined as a continuous frequency modulation in the spectrogram of at least 8 milliseconds in duration with a surrounding silent period of at least 16 milliseconds. In addition, a song is defined as a series of syllables with a surrounding silent period of at least 10 seconds.

### ICMS

Intra-cortical micro-stimulation was performed on head-restrained awake male mice to map cortical coordinates to complex behaviors (Clark, Armstrong, and Moore 2011; Gooler and O’Neill 1987; Pronichev and Lenkov 1998; Weiner et al. 2016). All mice were group housed and naive to females after weaning at P21. A broken pipette tip was lowered into the brain with a micromanipulator. Each site was stimulated between 10-100 times with a current stimulator of World Precision Instruments (WPI, FL). The stimuli consisted of either a 50ms (short) or 200-500ms (long) duration train of 333 Hz stimulation varying between positive and negative currents (biphasic square wave) between 10-100 µA. The large stimulus amplitudes are similar to other ICMS studies looking at complex behaviors (Bennett, Maier, and Brecht 2019; Graziano, Taylor, and Moore 2002)

### Fiber Photometry

We used multi-mode fibers (FT-200-URT, Thorlabs, Grünberg, Germany) with a diameter of 200 μm and a numerical aperture (NA) of 0.22. After removing the cladding from the fiber tip, the tip of the fiber was inserted into the brain and subsequently covered with dental cement. Low intensity stimulation (typically < 0.1mW at the tip of the optical fiber) was delivered by a 20 mW solid state laser (Sapphire, 488 nm, Coherent, Dieburg, Germany) for excitation of the GCaMP6s indicator and recording of Ca^2+^-dependent fluorescence was collected through the same fiber to an avalanche photodiode (APD, S5343, Hamamatsu Photonics, Herrsching, Germany). The resulting voltage signal was oversampled at 50Khz and stored for offline analysis.

### Optical Stimulation

*Guillardia* theta anion-conducting channelrhodopsin (GtACR2) was injected as described on a CamKII promotor in the AAV5 viral serotype (Malyshev et al. 2017, 2). Fiber optical cannulas were implanted as above into ACC. After recovery, male mice interacted with female mice in a pre-light baseline session. ACC neurons were suppressed in the second session after two days by optical stimuli from laser illumination (1.5-3 mW/mm^2^ using a 473-nm laser; Changchun New Industries Optoelectronics, Changchun, China). 50 Stimuli at 20Hz for a duration of 10 seconds were given semi-randomly, ensuring that about every 4 minutes at least 10 stimuli occurred. Short pulses of stimuli were given before the session began to calibrate the laser power to below the threshold for movement initiation.

### ICMS, Fiber Photometry, and Optical Stimulation Data Analysis

All analysis was conducted offline using custom code written in MatLab 9.9 (Mathworks). In ICMS experiments, each audio file was manually reviewed at least twice to identify any USVs. In fiber photometry experiments, relative fluorescence changes Δf/f = (f−f0)/f0 were calculated as Ca^2+^signals relative to baseline, where the baseline fluorescence was determined as the 10th percentile of the signal. For optical stimulation and fiber photometry experiments, the USV parsing was as described above.

### Neuropixel recording and analysis

The neuropixel probe is a high density and high signal-to-noise extracellular probe (Jun et al., 2017). The probe was soldered to short the reference pad to ground and ground was connected to a pin in the head-post. The probe was held on a custom metal-bar attached to a Luigs and Neumann (Ratingen, Germany) micromanipulator. Before recording, mice were habituated to head-restraint using their home-cage running wheel in the experimental setup. A brief craniotomy surgery was performed as described above to achieve access to cortex. The craniotomy was continuously filled with artificial cerebrospinal fluid (aCSF) and sealed between experiments. We recorded signals in 11 experiments from 3 mice in the left hemisphere ACC and M2 at a maximum depth between 2000-2500 μm. Probes were allowed to settle for about 10 minutes while probe gain calibration occurred. All recordings were made using SpikeGLX software. All data were preprocessed using the common average reference (CAR) by subtracting the common median across all channels (Ludwig et al., 2009). Single unit channels were selected by manual inspection. In the case that a single unit was on multiple channels, the channel with the largest spike waveform was chosen. Subsequently, spikes were detected as threshold crossings greater than 10 standards deviations from the mean.

### Video Analysis

Video was recorded at 30 frames per second concurrently with Neuropixel and sound data using a blue LED as a synchronization signal. Off-line, individual videos were manually cut and behavioral events were manually detected with start and end frames determining social exposure times.

### Statistical Analysis

Statistical significance was assessed using the unpaired Student’s two-sided t-test unless otherwise specified. Significant level was marked as *p < 0.05, **p<0.01, and ***p < 0.001. All data were reported as mean ± S.E.M. unless otherwise specified.

### Viral vector list

AAV9-CAG:: GCaMP6S.WPRE.SV40 (Penn Vector Core, titer 3.28 × 10^13^, 500 nl per site, LOT CS0775) AAV5-CK2:: GtACR2.Fred.kv.2.1 (ELSC Vector Core facility, titer 1 × 10^11^, 500 nl per site) AAVdj-CMV:: eGFP (ELSC Vector Core facility, titer 5 × 10^12^, 300 nl per site)

## Acknowledgements

This research has been supported by the Gatsby charitable foundation and a grant by the Israel Science Foundation (1024/17). B.W. has been supported by a fellowship of the Edmond and Lily Safra Center for Brain Science. We thank Shai Netser, Yoram Ben-Shaul, and David Omer for their insightful comments on the manuscript. We also thank Ofer Yizhar for providing the GtACR2 plasmids and Maya Groysman for vectors production.

## The authors declare no competing financial interests

### Author contributions

M.L and B.W. designed the experiments. B.W. performed all experiments, and analysis. M.L. and B.W. wrote the manuscript.

## Supp Figure

**Figure 4S:**
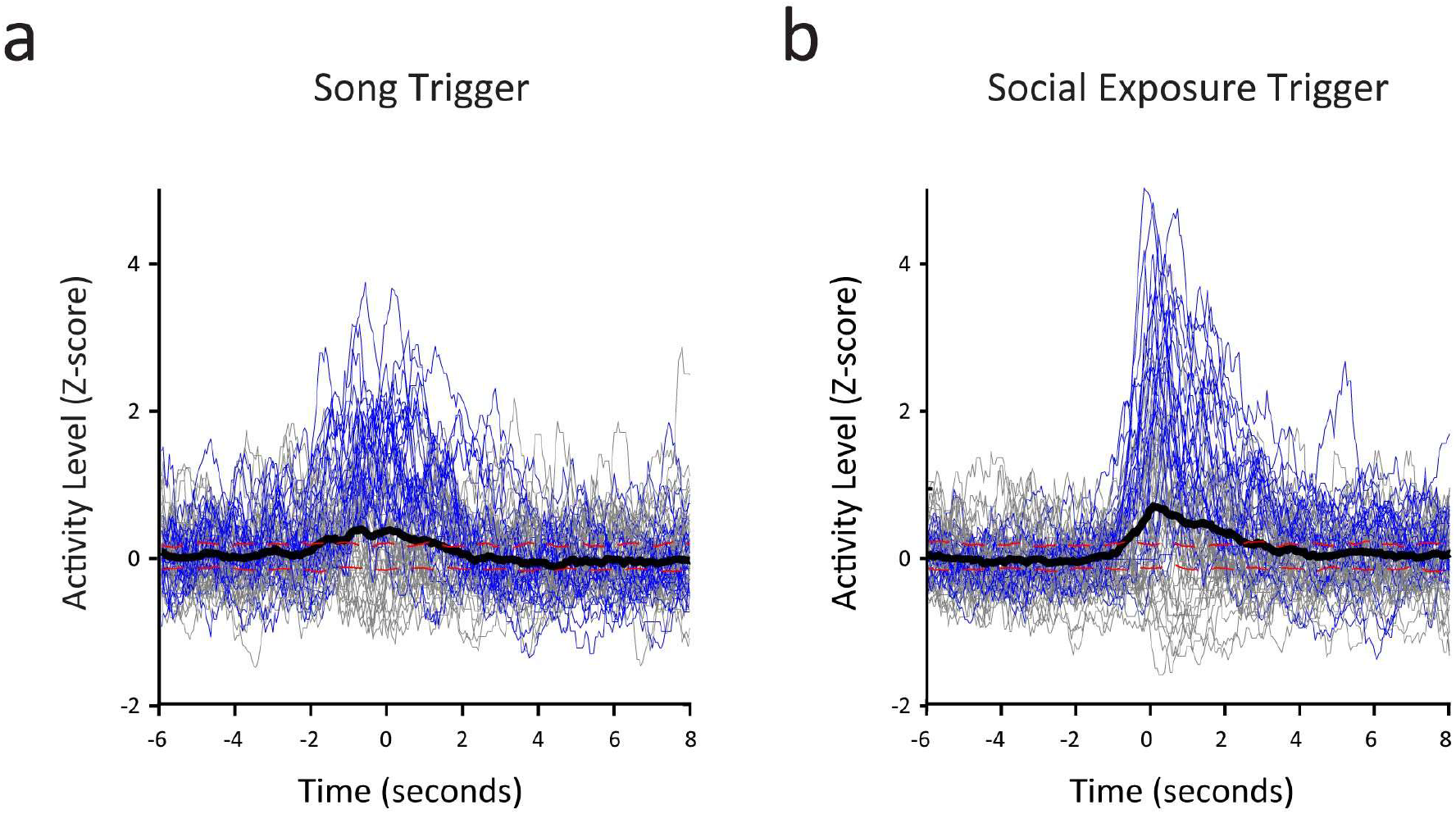
Averaging over all responsive neurons reveals a social exposure and song initiation response. **A**. Averaging all 130 responsive single units (black line) on social exposure. The result is similar, but weaker, than seen with only responsive neurons. Red dotted lines represent ±2 SD of a permutation test, while gray lines represent individual single units. Blue lines represent responding units (34/130). **B**. Same, but averaging at time zero on song initiations.

**Figure 6S:**
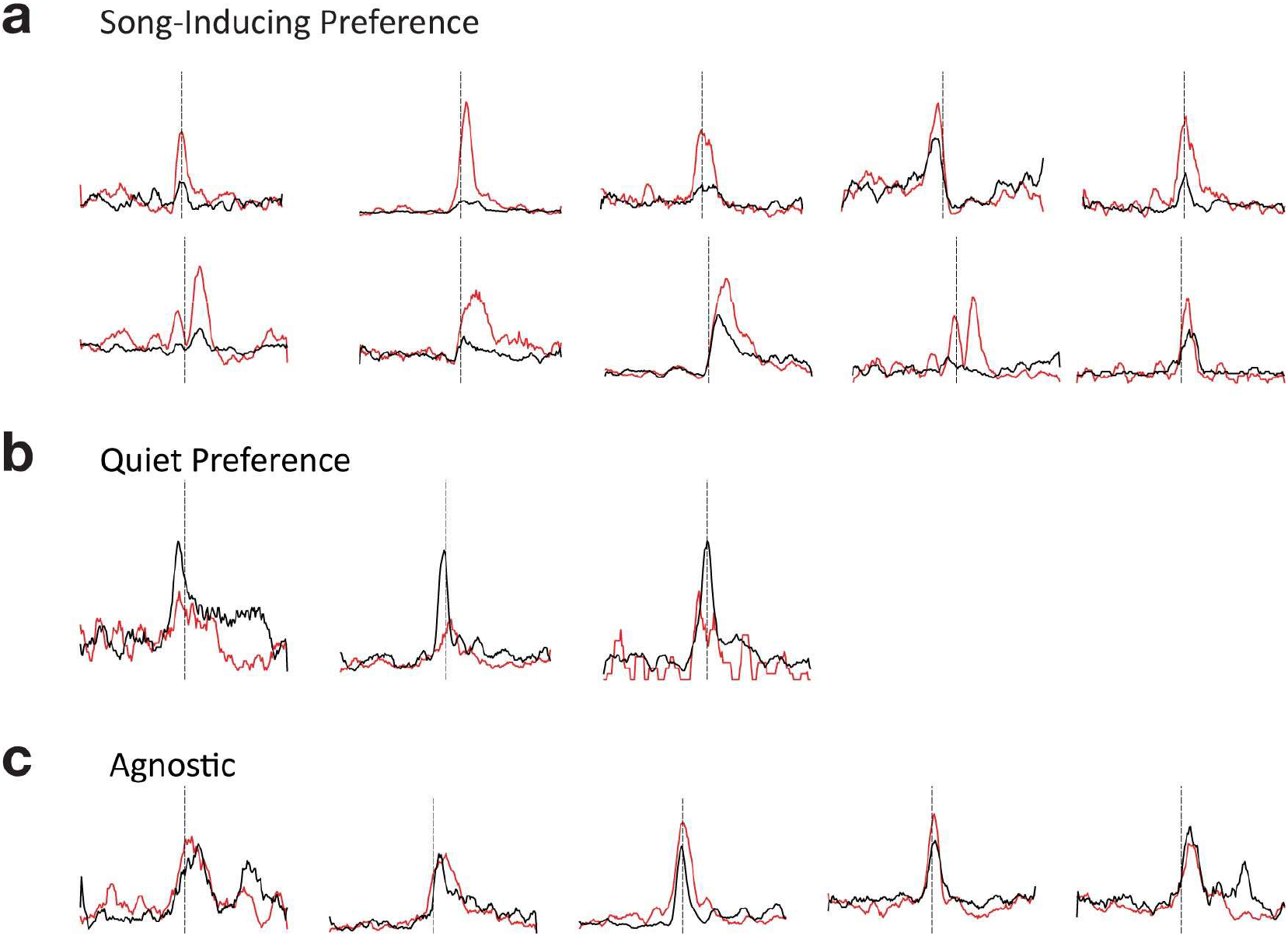
Single units were categorized by their response to social exposure into 3 groups: **A**. Song-Inducing, **B**. Quiet, or **C**. Agnostic. Most single units prefer the Song-Inducing category.

**Table S1:** Microstimulation ultrasonic vocalization parameters.

|  |  |
| --- | --- |
| Duration(ms) | 30.50 +- 23.62 |
| Freq Start (kHz) | 84.17 +- 15.75 |
| Freq Mean (kHz) | 81.08 +- 14.08 |
| Freq Max (kHz) | 90.83 +- 16.86 |
| Freq Bandwidth (kHz) | 18.18 +- 14.34 |

## Notes

### Competing Interest Statement

The authors have declared no competing interest.

